# The sex-limited effects of mutations in the EGFR and TGF-β signaling pathways on shape and size sexual dimorphism and allometry in the *Drosophila* wing

**DOI:** 10.1101/037630

**Authors:** Nicholas D. Testa, Ian Dworkin

## Abstract

Much of the morphological diversity in nature-including among sexes within a species-is a direct consequence of variation in size and shape. However, disentangling variation in sexual dimorphism for both shape (SShD), size (SSD) and their relationship with one another remains complex. Understanding how genetic variation influences both size and shape together, and how this in turn influences SSD and SShD is challenging. In this study we utilize *Drosophila* wing size and shape as a model system to investigate how mutations influence size and shape as modulated by sex. Previous work has demonstrated that mutations in Epidermal Growth Factor Receptor (EGFR) and Transforming Growth Factor - ß (TGF-ß) signaling components can influence both wing size and shape. In this study we re-analyze this data to specifically address how they impact the relationship between size and shape in a sex-specific manner, in turn altering the pattern of sexual dimorphism. While most mutations influence shape overall, only a subset have a genotypic specific effect that influences SShD. Furthermore, while we observe sex-specific patterns of allometric shape variation, the effects of most mutations on allometry tend to be small. We discuss this within the context of using mutational analysis to understand sexual size and shape dimorphism.

## Introduction

In spite of our wealth of knowledge about the natural world, biologists continue to be fascinated by the prevalence of sexual dimorphism. Where sexual dimorphism is often found, it is most often subtle, despite important exceptions of sex-limited characteristics (Bonduriansky & Day 2003), or traits that are highly exaggerated in one sex, but not the other (Lavine et al. 2015). This is particularly evident for morphological traits that demonstrate sexual size (SSD) or sexual shape (SShD) dimorphism (Kijimoto et al. 2012). Within evolutionary biology, explanations for sexual dimorphism have focused on a number of mechanisms that are likely responsible for the origin and maintenance of sexual dimorphism (Reeve & Fairbairn 2001; Allen et al. 2011; Bonduriansky & Chenoweth 2009; Mank 2009; Cox & Calsbeek 2010; Hedrick & Temeles 1989; Shine 1989; Fairbairn & Blanckenhorn 2007) including sexual conflict, differences among the sexes in the variance of reproductive success leading to sexual selection (Fairbairn 2005), and sex specific aspects of natural selection (Preziosi & Fairbairn 2000; Ferguson & Fairbairn 2000). Despite this, our understanding of the genetic mechanisms that contribute to variation in sexual shape and size dimorphism is still lacking (Mank 2009; Blanckenhorn et al. 2007; Fairbairn & Roff 2006; Fairbairn 1990).

There is considerable experimental evidence demonstrating that patterns of SSD and SShD can be altered by influencing the condition of individuals (Bonduriansky & Chenoweth 2009; Bonduriansky 2007). There has unfortunately been less success on directly experimentally evolving consistent changes SSD or SShD, with some notable exceptions where dimorphism evolved in response to selection on fecundity (Reeve & Fairbairn 1999) or due to experimental manipulation in the degree of sexual conflict (Prasad et al. 2007). There are even fewer instances where experimental evolution has been able to alter existing size/shape (allometry) relationships (Bolstad et al. 2015).

Despite previous difficulties with directly selecting for SSD or SShD, we still find evidence for genetic variation in SSD within a number of species (David et al. 2003; Merila et al. 2011). Several studies have utilized induced mutations (Carreira et al. 2011) or defined genomic deletions to examine patterns of SSD (Takahashi & Blanckenhorn 2015). They find that, in general, mutations tend to attenuate differences in SSD and sexual developmental timing difference. Interestingly, while ˜50% of the random insertion mutations influenced size and shape, only half of those were consistent between males and females, suggesting considerable sex limitation of the mutational effects (Carreira et al. 2011).

With respect to the influence of mutations on sexual dimorphism, one important consideration is whether the mutations themselves are directly influencing aspects of sexual dimorphism. Alternatively, mutations may be influencing size and shape of the organism, but are modulated in a sex-limiting fashion. Arguably, it is difficult to distinguish between these possibilities, although for the purposes of this study, we consider a mutation to be modulated by the influence of sex if it influences size or shape as well as having an additional influence on sex (i.e. a sex-by-genotype interaction). The extent to which such mutations influence SSD and SShD remains poorly understood.

To address these questions, we examined the influence of characterized induced mutations that influence two signaling pathways important for wing development, Epidermal Growth Factor Receptor (EGFR), and Transforming Growth Factor - β, (TGF-β). The *Drosophila* wing is an excellent model for the study of SSD and SShD. First, as it is a premiere model system for the study of development, and as such a great deal is known and understood about the mechanisms governing overall growth and patterning (García-Bellido et al. 1994; Weinkove et al. 1999; Day & Lawrence 2000; Weatherbee et al. 1998). Additionally, *Drosophila melanogaster* and closely related species have a strong pattern of sexual size dimorphism for many traits (and overall body size), with wing size demonstrating some of the greatest degree of overall dimorphism (Testa et al. 2013; Abbott et al. 2010; Gidaszewski et al. 2009). There is extensive variation for size and shape within and between *Drosophila* species, and for the extent of SSD and SShD as well (Gidaszewski et al. 2009). Importantly, the mutational target size for wing shape (Weber 2005) is high (˜15% of the genome), thus providing plenty of opportunity for mutations to influencing shape, and potentially those modulated by sex.

In this study we utilize a previously published data set that examine the influence of 42 mutations in the EGFR and TGF-ß signaling pathways when examined in a heterozygous state. We re-analyze this data set to examine the extent to which the mutations have sex-limited phenotypic effects that influence SSD or SShD. Furthermore we examine how patterns of allometric variation between size and shape are altered by both sex and wild type genetic background of the mutations. Despite most mutations having substantial phenotypic effects on either size, shape or both, only a small subset of them appear to have their effects modulated by sex, with respect to both direction and magnitude of effects. Furthermore, we demonstrate that the allometric relationship between size and shape is only subtly influenced by sex and genetic background for these alleles. We discuss these results within the context of sex-limited effects of mutations and their influence on SSD and SShD, and how to interpret allometric relationships between size and shape in *Drosophila.*

## Materials and Methods

### Provenance of Samples

The data used for this study was originally published by (Dworkin & Gibson 2006). We compared wings from flies across several treatment groups, including: sex, wild type genetic background (Oregon-R and Samarkand), progenitor line and genotype (mutant vs. wild type allele). Fifty different p-element insertion lines, each marked with *w*^+^, were introgressed into two common wild type backgrounds (Samarkand and Oregon-R), were used along with their respective controls. All wing data, in the form of landmarks, were collected from digital images, as detailed in Dworkin and Gibson (2006). For a more detailed description on the source of these strains and the experimental design, please refer to Dworkin and Gibson (2006).

Insertional mutations were selected from the Bloomington Stock Center and subsequently introgressed into two wild-type lab strains, Samarkand (Sam) and Oregon-R (Ore). Introgressions were performed by repeated backcrossing of females bearing the insertion to males of Sam and Ore-R. Females from replicate vials within each generation were pooled for the subsequent generation of backcrossing. Since both backgrounds contain a copy of the mini-white transgene, eye color for all flies lacking p-elements was white. Selection was therefore based entirely on the presence of the eye color marker, precluding unwitting selection for wing phenotypes. While the introgression procedure (14 generations of backcrossing) should make the genome of the mutant largely identical to that of the isogenic wild types, some allelic variation in linkage disequilibrium with the insertional element may remain. All experimental comparisons of mutant individuals were therefore made with wild-type siblings from a given cross and should share any remaining segregating alleles unlinked to the p-element. We separated mutants and their wild-type siblings by their corresponding mutant “line” number (supplementary Table 1) to avoid these and potential “vial effects”. All crosses were performed using standard media, in a 25°C incubator on a 12/12-hr light/dark cycle.

Two vials for each line were set up carefully to result in low to moderate larval density. The temperature of the incubator was monitored cautiously for fluctuations, and vial position was randomized daily to reduce any edge effects. After eclosion and sclerotization, flies from each cross were then separated into mutant and wild type individuals—those with and without the p-element-induced mutations, respectively—based on eye color and stored in 70% ethanol. A single wing from each fly was dissected and mounted in glycerol (see supplementary table 1B for sample sizes). Images of the wings were captured using a SPOT camera mounted on a Nikon Eclipse microscope. Landmarks (as shown in Figure 1) were digitized using tpsDig (v. 1.39, Rohlf 2003) software.

**Table 1.** Summary table of significant effects by background among mutants. All significant values are taken om Figures 1-4. In the case of vector correlations, 80% was chosen arbitrarily to represent only a small bset of mutants of large effect.

| **Mutant** | **Allele** | **Pathway** | **$\Delta$ SSD** | **$\Delta$ SShD** | **$\Delta$ Vector Correlation** | **$\Delta$ Allometry** |
| --- | --- | --- | --- | --- | --- | --- |
| *aos* | W11 | Egfr |  |  |  |  |
| *omb* | md653 | TGF-β | Ore | Sam, Ore | Sam, Ore | |
| *cv-2* | 225-3 | TGF-β | | | | |
| *GAP1* | mip-w[+] | Egfr |  |  |  |  |
| *ksr* | J5E2 | Egfr |  |  |  | Sam |
| *dad* | J1E4 | TGF-β | Ore | Ore | | |
| *drk* | k02401 | Egfr |  | Ore |  |  |
| *bs/DSRF* | k07909 | Egfr |  | Ore |  | Sam |
| *s* | k09530 | Egfr |  |  |  | Sam |
| *spi* | s3547 | Egfr |  |  |  |  |
| *mad* | k00237 | TGF-β | | | | Ore |
| *ed* | k01102 | Egfr |  |  |  | Sam |
| *tsh* | A3-2-66 | TGF-β | Sam | | | |
| *cos* | k16101 | Hh |  |  |  |  |
| *tkv* | k19713 | TGF-β | | | | |
| *babo* | k16912 | TGF-β/Hh | | | | |
| *trl* | S2325 | TGF-β | | | | |
| *rho-AP* | BG00314 | ? | Sam |  | Sam |  |
| *pka-C1* | BG02142 | Hh |  |  |  |  |
| *sbb* | BG01610 | TGF-β | | | | |
| *psq* | kg00811 | Egfr | Ore |  |  | Ore |
| *osa* | kg03117 | Chromatin Remodeling |  |  | Sam |  |
| *rasGAP* | kg02382 | Egfr |  |  |  |  |
| *pnt* | kg04968 | Egfr | Ore | Ore | Ore |  |
| *drk* | k02401 | Egfr |  |  |  |  |
| *cbl* | kg03080 | Egfr |  |  |  |  |
| *mam* | kg02641 | N/Egfr |  |  |  |  |
| *rho-6* | kg05638 | Egfr |  | Ore | Sam, Ore |  |
| *dpp* | kg04600 | TGF-β | | | Ore | Sam |
| *pka-C3* | kg00222 | Hh |  |  | Sam | Sam |
| *p38b* | kg01337 | TGF-β/Egfr | | | | |
| *tkv* | kg01923 | TGF-β | | | | |
| *wmd* | kg07581 | Unknown |  |  |  |  |
| *mad* | kg00581 | TGF-β | | | | |
| *ast* | kg07563 | Egfr |  |  |  |  |
| *dpp* | kg08191 | TGF-β | | | | |
| *rho1* | kg01774 | Egfr? |  |  |  | Sam |
| *sax* | kg07525 | TGF-β | | | | |
| *sax\** | sax4 | TGF-β | Sam | Sam | Sam | |
| *egfr* | k05115 | Egfr |  | Sam, Ore | Ore | Sam |
| *src42A* | kg02515 | Egfr |  |  |  | Ore |
| *rho/stet* | kg07115 | Egfr |  |  |  |  |

**Figure 1.**
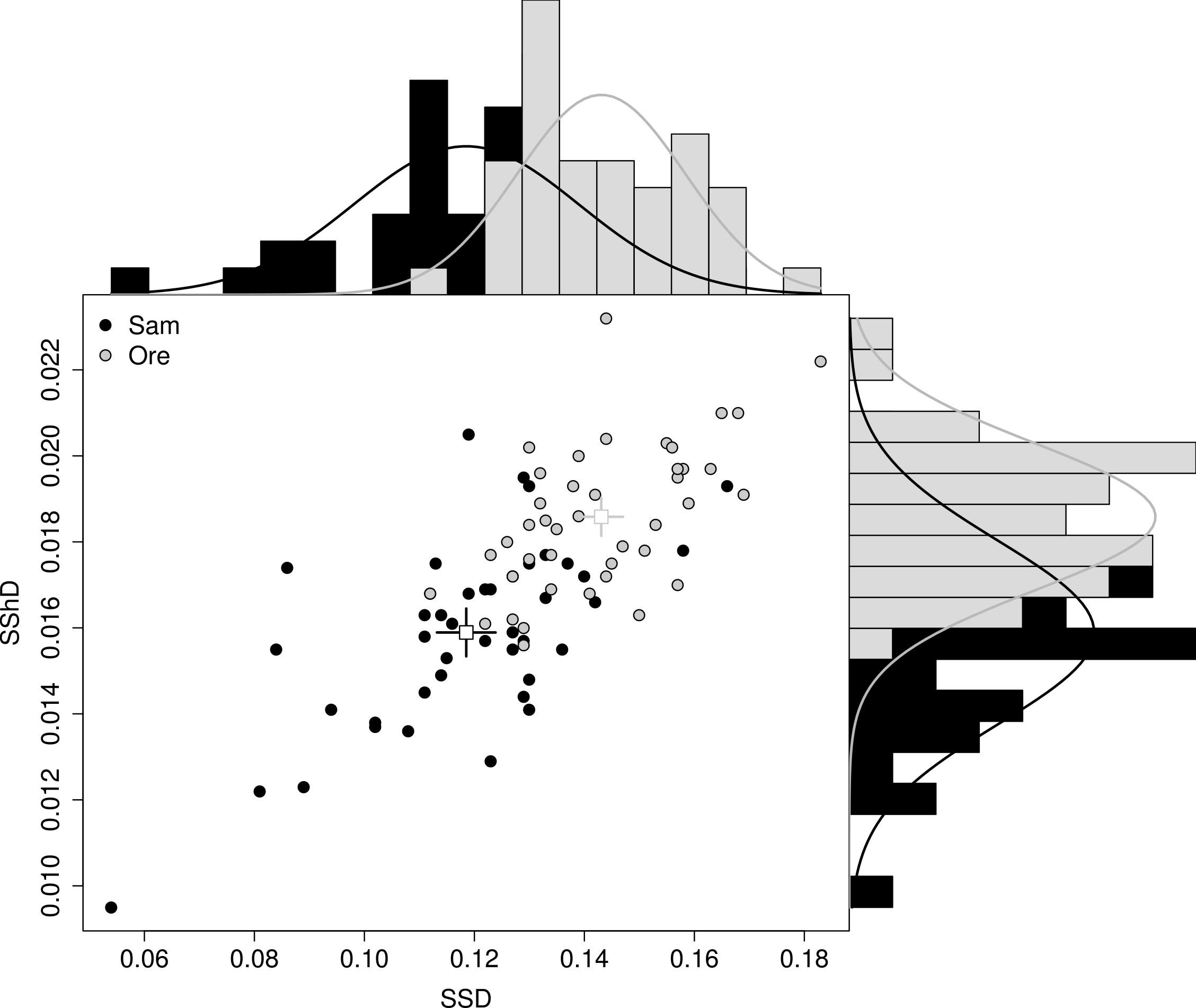
Natural variation in SSD and SShD for two wild type strains. A) SSD and SShD in both wild type background strains are represented three ways: scatterplot (center), SSD histogram with density curve (x-axis, top) and SShD histogram with density curve (y-axis, right). Data points represent mean value for wild-type siblings of each heterozygous mutant cross from a given vial. The Samarkand wild type background has a wider range of SSD, encompassing the low end of the spectrum, whereas Oregon-R tends to be more consistently large in SSD. B) Average direction of SShD in a typical Samarkand (left) and Oregon-R (right) wild type wing. Landmark coordinates are mapped onto a typical wing to demonstrate shape. Arrows represent the vector of shape change (magnified 5x) from female to male wing shapes.

Our analysis necessitated that there be flies from each representative treatment group; those lines with flies missing (e.g. from one background or sex) were left out of the analysis. Of the original 50, 42 lines were ultimately used.

### Analysis of Sexual Size Dimorphism (SSD)

Centroid size (i.e., the square root of the sum of the squared distances from each landmark to the centroid of the configuration) was used as the size variable in our analyses. Individual size values for male and female within each line and background were taken from the coefficients of a linear model where centroid size was modeled as a function of genotype, sex and their interaction.

SSD was then calculated based on a common index, wherein the dimorphism is represented as the proportion of female size to male size (Lovich & Gibbons 1992; Smith 1999):

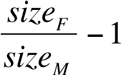

The resulting index represents the relative size difference between males and females where 0 indicates a complete lack of dimorphism and 1 indicates that females are 100% larger than males. Negative values represent male-biased dimorphism.

### Analysis of Sexual Shape Dimorphism (SShD)

Generalized Procrustes Analysis (GPA) was used to super-impose landmark configurations after correcting for position and scaling each configuration by its centroid size. This procedure removes non-shape variation from the data—size, orientation and position. From the nine two dimensional landmarks, we are left with 14 dimensions of variation, and thus applied a Principal Components Analysis (PCA) to the Procrustes coordinates (i.e., the shape coordinates after GPA) and the first 14 PC scores were used as shape variables in subsequent shape analyses.

Two different shape scores were used in this study: one to examine sexual shape dimorphism and one to assess the strength of the allometric relationship of shape on size. First, SShD was estimated using the tangent approximation for Procrustes distance (i.e. Euclidian distance) between the average of male and female wing shape for a given treatment. Additionally, we calculated shape scores from the multivariate regression of shape onto size based on Drake and Klingenberg (2008). Specifically we projected the observed shape data onto the (unit) vector of regression coefficients from the aforementioned multivariate regression. We used these shape scores and regressed them onto centroid size to approximate allometric relationships. Confidence intervals for SSD and SShD as well as allometric coefficients were generated with random non-parametric bootstraps, using 1000 iterations.

All significance testing for the analyses involving shape data was done with Randomized Residual Permutation Procedure (RRPP) as implemented in the *geomorph* library in R. (Collyer et al. 2015). This method differs from the analyses in the original paper in two important ways. First, the linear model is based upon Procrustes distances, and second the resampling procedure more easily enables inferences within nested models (Collyer et al. 2015) with interaction terms. Specifically, this approach samples (without replacement), the residuals from the “simple” model under comparison, adding these to fitted values, and refitting under the “complex” model. We used the following models to assess the difference in shape dimorphism for each line and wild type background:

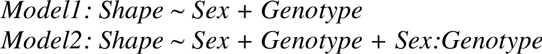

We then performed such analysis for increasing degrees of interactions for the influence of sex, genotype, genetic background and size (for models of shape variables).

SShD was calculated with one of two methods: the advanced.procD.lm() function in the *geomorph* package (v.2.1.8) in R (v. 3.2.2) and standard Euclidean distances among treatment groups using the lm() function; both approaches yielded equivalent results. To evaluate the mean shape difference caused by sex, we used linear models based upon Procrustes distance (with RRPP) to compare models where sex is and is not a predictor of shape using the procD.lm and advanced.procD.lm functions in *geomorph.* These analyses were randomized (by individual) and repeated 1000 times per treatment group to assess whether the magnitude of effect was greater than expected by chance.

Despite having separate and independent “control” (wild type) lineages for each cross (to control for any potential vial effects or residual segregating variation), we utilized a sequential Bonferroni correction to maintain our “experiment-wide” nominal alpha of 0.05. Given the large number of comparisons being made, it is likely that this will yield extremely conservative results, and we expect this underestimates the number of mutations that influence sexual dimorphism or mutational effects of allometry of shape on size.

### Vector Correlations

While the above linear model assesses the magnitude of the effects, for shape it is also important to examine the direction of effects. Specifically, whether the mutations influenced the direction of SShD. To examine this, the vector of SShD was calculated within each genotypic group (wild type VS. mutant). We then estimated the vector correlation between the vectors of SShD for the wild type and mutant as follows:

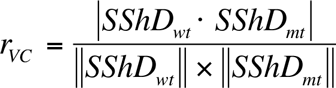

Where the SShD for each genotype is equal to difference between the female and male vectors within each genotype. We used the absolute value of the numerator to avoid arbitrary sign changes. The denominator consists of the product of the length (norm) of each vector. As with a Pearson correlation coefficient, a value of 0 corresponds to no correlation, while a value of 1 means that each vector is pointing in the same direction (even if they differ in magnitude). Approximate 95% confidence intervals were generated using a non-parametric bootstrap of the data for each line (The alpha used for the 95% CIs were not adjusted for the number of mutant alleles tested).

### Statistical Analysis

All statistical analyses were conducted using R statistical software (version 3.2.2). Significance testing (specifically those involving RRPP) was conducted using functions within the *geomorph* package (v. 2.1.8) and with custom functions. All error bars are 95% Confidence intervals generated by non-parametric bootstraps. All scripts including custom functions are available on github (https://github.com/DworkinLab/TestaDworkin2016DGE).

## Results

### Different wild type strains vary for Sexual Size Dimorphism (SSD) and Sexual Shape Dimorphism (SShD) in wing morphology, for both direction and magnitude

As each mutation was repeatedly backcrossed into two distinct wild type strains—Oregon-R (Ore) and Samarkand (Sam)—we first examined patterns of sexual size and shape dimorphism between these two strains.

We observed considerable, and highly significant, differences in both SSD and magnitude of SShD between the wild type strains (Fig.1A). Further, with respect to the vector of SShD, both wild-type backgrounds were somewhat divergent (Fig. 1B). The computed vector correlation for SShD between both backgrounds falls within the same range as those calculated for SShD by genotype (0.937, 95% CI 0.92, -0.95), suggesting only subtle changes in direction. Additionally, the allometric relationship between shape and size differs between the two wild type backgrounds. While Ore has a stronger overall slope than Sam, the magnitude of both males and females slopes are reversed by background; for females, shape has stronger association with size relative to males in Ore (F 0.113, 95% CI 0.105, 0.122; M 0.099, 95% CI 0.091, 0.107), whereas the opposite is true for Sam (F 0.105, 95% CI 0.097, 0.113; M 0.120, 95% CI 0.112, 0.129). These differences in size, shape and allometry are all significant based on the randomized resampling permutation procedure (see methods).

Despite tight control of experimental variables (food, temperature) we observed a surprising amount of residual environmental variation for SSD and SShD among each replicate of the two wild type lineages. In the design of the experiment, where for each mutation, within each background, wild-type controls were generated from the cross that shared the environment (vials) with their otherwise co-isogenic mutant sibling. As all of these offspring across the vials are genetically co-isogenic, and only differ in the subtle aspects of rearing environment across vials, this allows us to assess some aspects of how environmental variation influences SSD and SShD. As shown in Figure 1A, in addition to differences between the two wild type strains for SSD and SShD, there is also variation around the mean estimates for each. Since each data point in Figure 1A corresponds to each mutant’s wild-type siblings from a given cross, these points largely reflect variation among “vial” effects. Indeed, models based on Procrustes distance suggest that there are significant vial effects (P = 0.009) and vial by sex (P = 0.001) even within the background control populations, which are largely attributable to micro-environmental variation. This is somewhat surprising as external sources of variation such as food (all from a common batch) and rearing temperature (all vials reared in a common incubator, with daily rotation of vials to minimize edge effects) were highly controlled in the experiment. This suggests that the magnitude of SSD and SShD for wing form is influenced by subtle environmental changes, suggesting that high levels of replication to control for these factors is generally necessary.

### Despite many mutations having substantial effects on overall shape, a relatively small number influence SSD and SShD

As demonstrated in the original study (Dworkin & Gibson 2006) and confirmed here, the vast majority of mutations have a significant influence on shape when measured in the heterozygous state (supplementary table 1). Of the subset of 42 mutations used in the current study (from the original 50), all but 10 had a significant effect for genotype (most surviving even a conservative Bonferroni correction) using the Residual permutation (Collyer et al. 2015). Of those 10, most had significant genotype-by-background effects, consistent with the earlier study (despite a different underlying inferential approach). Despite this, only 18 of the mutations showed evidence for “significant” sex-limited genotypic effects (based on the sex-by-genotype effects), of which 2 survived sequential Bonferonni correction. Additionally, another 12 show evidence for significant effects of sex-by-genotype in combination with other factors in the model (size and/or background). Only one of these 12 survived correction for multiple comparisons. While inferences based on significance alone is quite limited (see below), these results suggest that only a small subset of mutations appear to have sex-limited influences on shape (Table 1).

To understand these results more fully, we next focused on the magnitudes of SShD and the SSD index, using non-parametric bootstraps to generate confidence intervals on our estimates. We performed the analyses separately for each wild type genetic background given that they can differ for both magnitude and direction of SShD. As shown in Fig.2, while several mutants show significant effects for either SSD, SShD or both in one or both of the backgrounds, the magnitudes of these effects are small, especially considering the relatively large amount of environmental variation in SSD and SShD observed within strains (Fig.1A). Interestingly, while only a few mutations showed evidence for an overall effect on size, these tend to have sex-limited effects (Fig.2).

**Figure 2.**
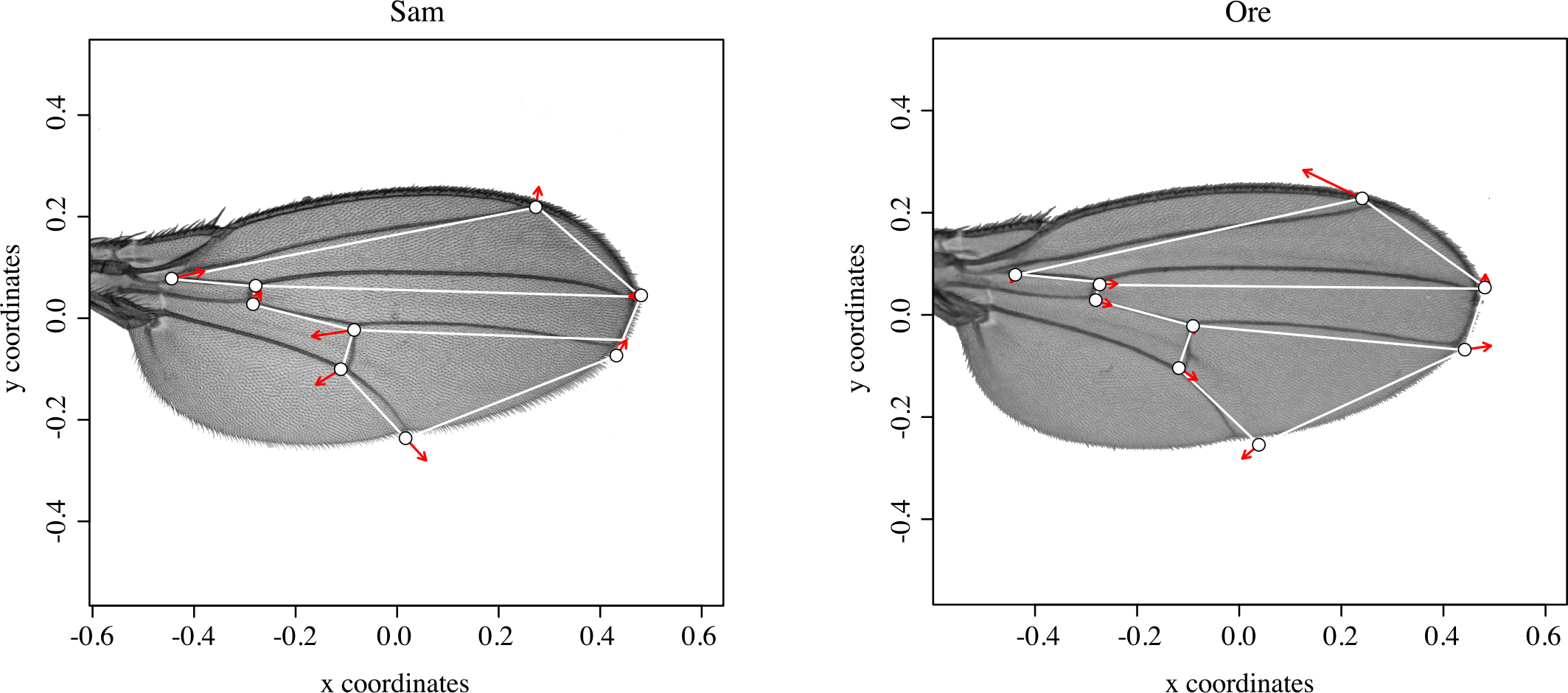
Magnitude of SSD and SShD for 42 mutants in Oregon-R (left) and Samarkand (right) wild type backgrounds. The effect of each mutant is mapped out in a size-and-shape dimorphism space. Genotypic means for each mutant are indicated by point style and connected by a solid line. SSD is plotted on each x-axis for all plots and SShD is displayed on the y-axis. The plots above display the entire range of variation observed, while those below display only the area with the highest density of points. Lines with significant sex-by-genotype effects are highlighted as follows: effect on both size and shape, shape only and size only. Only significant genes (after sequential Bonferroni correction) from the linear models are colored. Few mutations in this study alter sexual dimorphism of size or shape. In addition, the effect of mutations also appears to be highly background dependent, as only two lines, *Omb* and *Egfr,* were consistent in both backgrounds. Error bars are 95% confidence intervals (unadjusted alpha). All gene names are displayed lower-case, regardless of dominance.

In addition to examining the magnitude of effects, we also examined the direction of effects, and whether the mutations substantially changed the direction of SShD relative to their co-isogenic wild type. As shown in Fig.3, the mutations examined in this study generally do not substantially influence the direction of SShD, with several notable exceptions such as the mutation in the *Omb* gene, as well as more subtle effects from mutations such as *sax, pnt, drk* (among others). Even when the bootstrap confidence intervals do not approach 1, the estimated vector correlation are still generally greater than ˜0.9, suggesting only modest changes in the direction of SShD.

**Figure 3.**
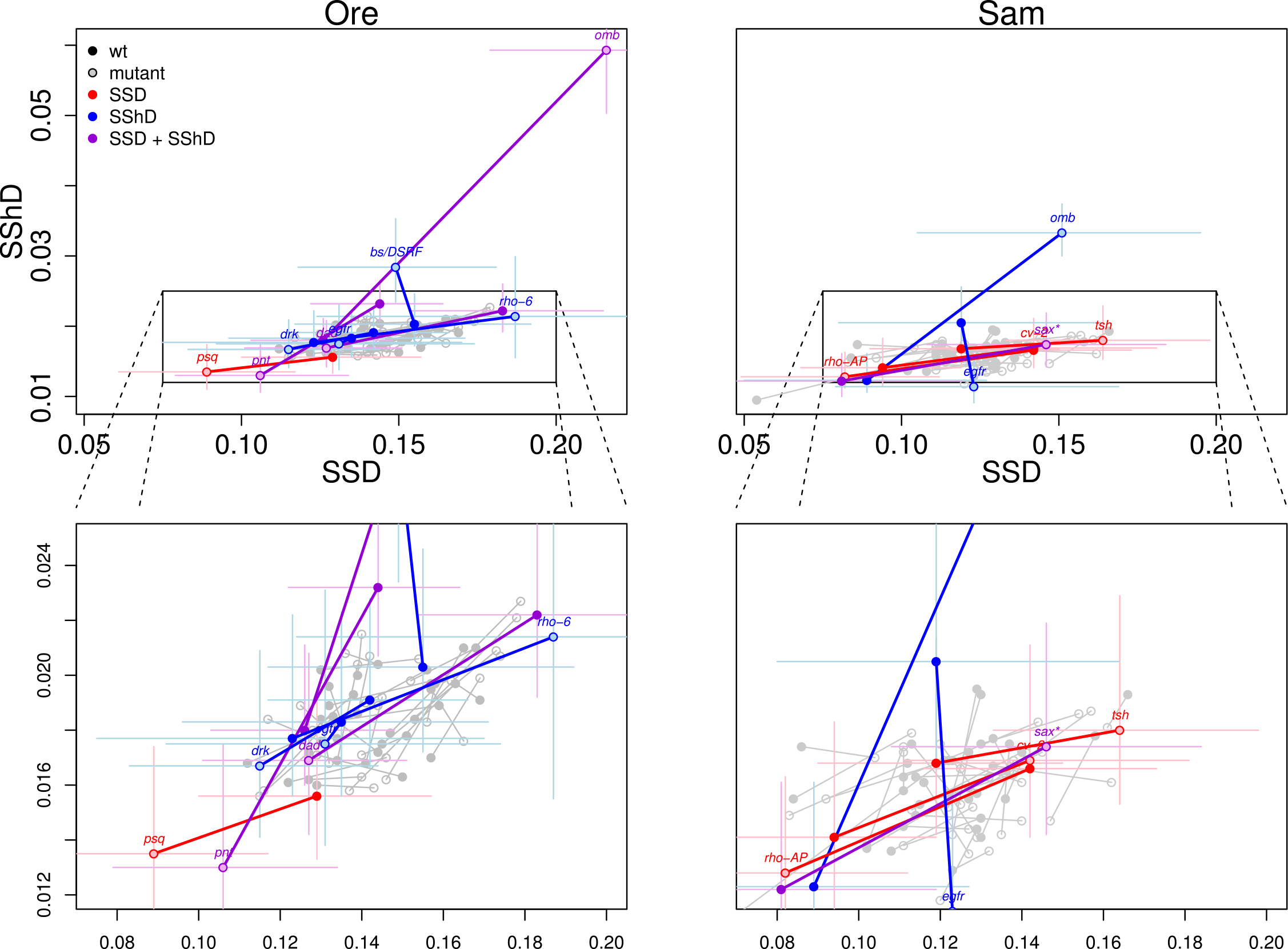
Vector correlations to assess similarity of direction for sexual shape dimorphism (mutant VS. wild type) by background. While genetic background appears to have little effect on the direction of SShD for most mutations, several stand out with more divergent directions of SShD. Those mutations with large background effects are also notable for their large effect on size and/or shape. Error bars are 95% confidence intervals (unadjusted alpha).

### Mutations do not substantially alter directions of SShD, nor patterns of allometry

One important aspect of assessing variation in shape, and in particular in situations where there is either (or both) SSD or SShD, is to account for the allometric effects of size on shape when computing the magnitude and direction of SShD. One important approach is to assume a common allometric relationship between size and shape across the sexes (after adjusting for mean differences in size and shape), and regressing out the effects of size. Then using either the residuals or predicted values of shape (after accounting for size) to compute an “allometry corrected” measure of SShD (Gidaszewski et al. 2009). To utilize such an approach requires that the assumption of a common allometric relationship be valid, as has been observed across *Drosophila* species for the wing shape and size relationship (Gidaszewski et al. 2009).

Prior to computing the allometry-corrected measure we examined this assumption among the mutations used in this study. Of the 42 independent mutations (with their independent controls), 13 had a significant interaction of sex-by-size on the influence of shape (with 3 surviving the sequential Bonferroni correction). Another 8 of them had a sex-by-size interaction imbedded within a higher-order interaction term. Despite this the overall magnitudes of effects and directions of allometric relationships appear to be highly similar, with a few important exceptions (Fig.4). Thus it is unclear whether using an allometry-free correction is warranted within the context of this study. It is worth noting that making the assumption of a shared allometric relationship, and computing the allometry-corrected measure of SShD did not substantially alter our findings (Supplementary Figure 1; Supplementary Table 2).

**Figure 4.**
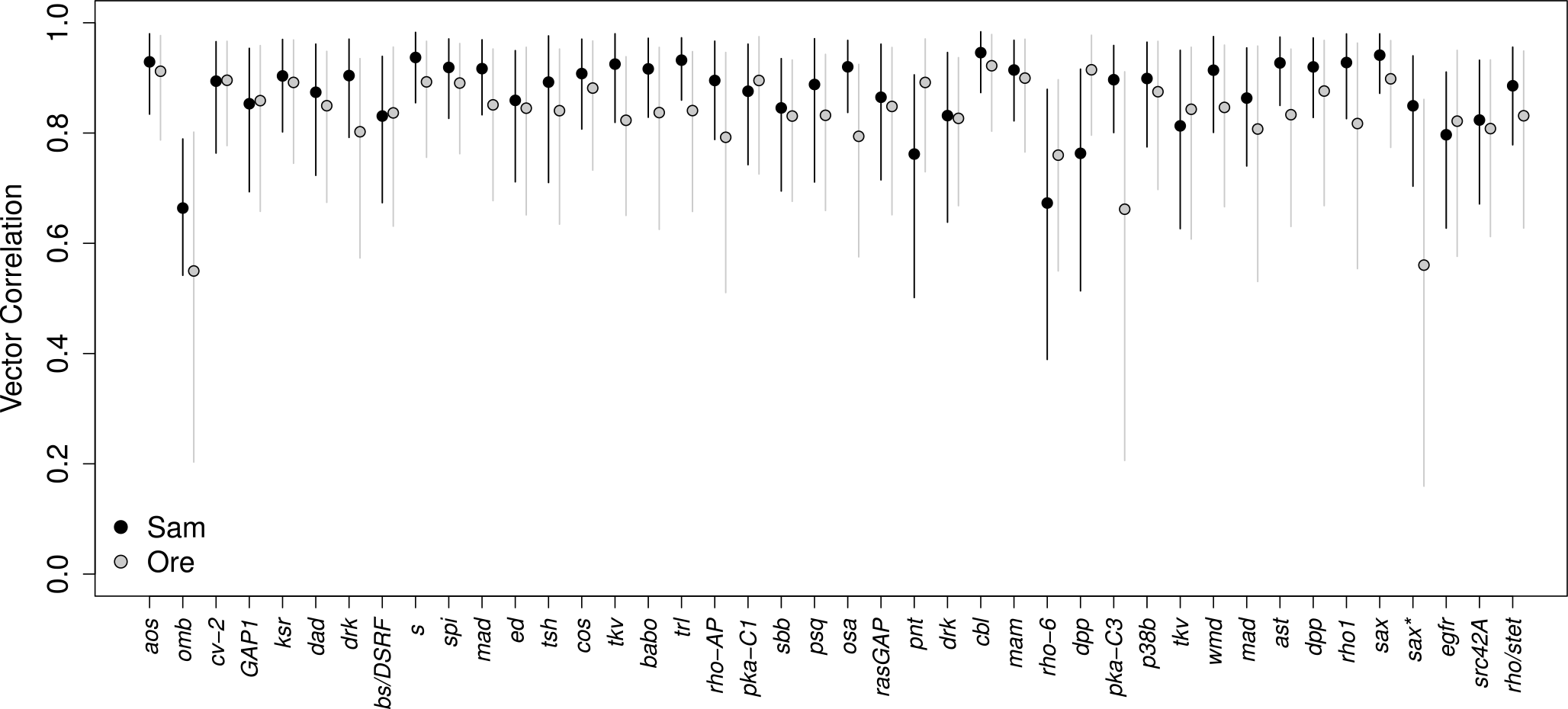
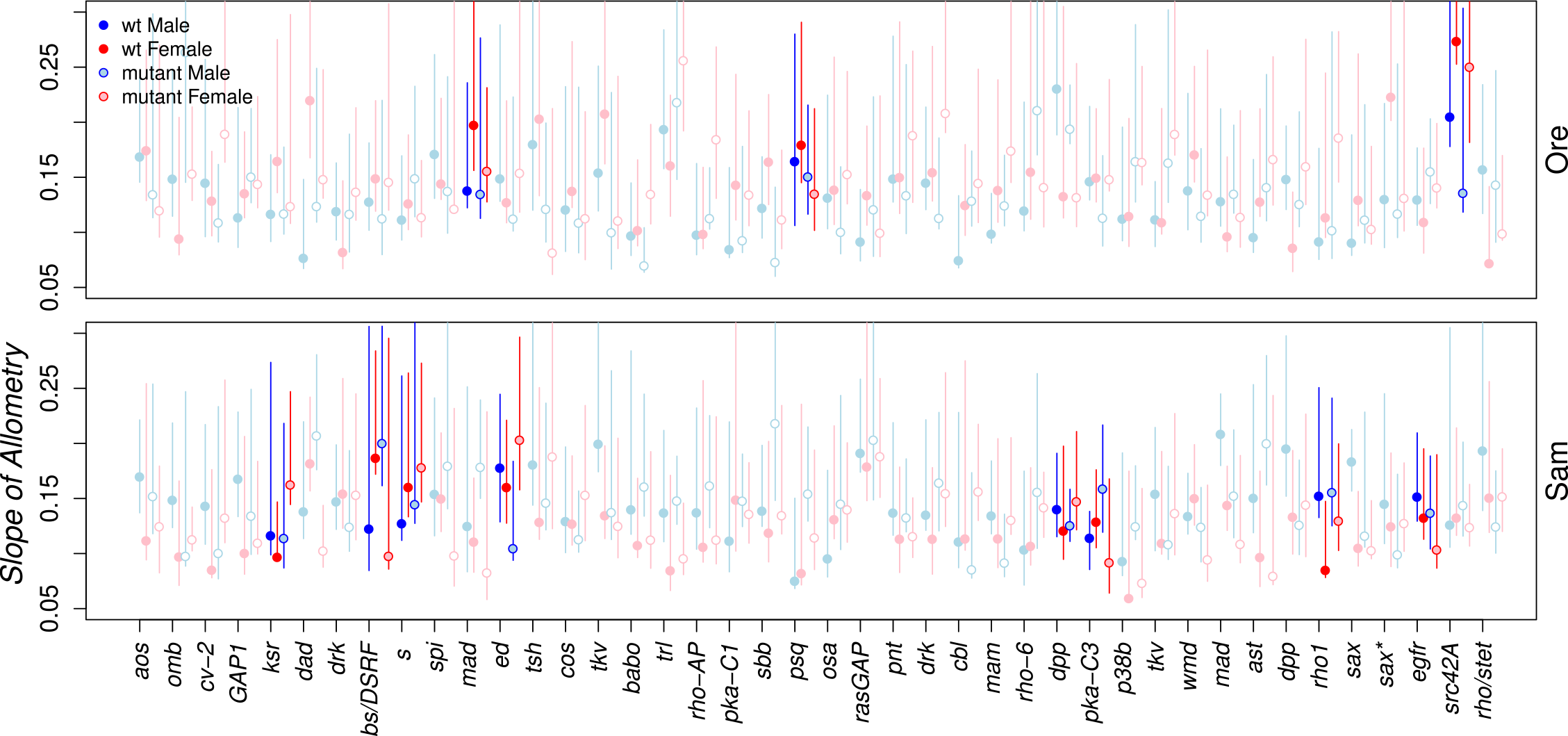
Variation in the magnitude of association between shape and size allometric coefficients among mutations in the Oregon-R (top) and Samarkand (bottom) wild type backgrounds. The “slope” of the allometric relationship for shape on size is displayed by sex and genotype. The magnitude of allometric effects appears to be relatively stable across strains, with few mutants substantially altering the wild type pattern of allometric co-variation. Individual lines whose mutants cause a significant sex-by-size interaction are represented dark in contrast to non-significant (faded) lines. Error bars are 95% confidence intervals (unadjusted alpha) for each individual treatment; significance is assessed based solely the interaction terms from the multivariate linear models.

### Discussion

While previously underappreciated, it is clear that mutations in genes in several growth factor pathways can act in a sex-specific manner. Of the 42 mutations analyzed, 12 had a significant sex-by-genotype interaction on size, shape or both (Fig.2). Only a few mutant alleles had the ability to affect the sexual dimorphism in allometry, the relationship between shape and size (Fig.4). Furthermore, nearly all of the mutants appear to act in a background-dependent manner, affecting shape or size in one genotype, but not the other (Figure 2).

Previous research has demonstrated the ability of growth pathways to respond to various perturbations, including: individual mutation (Palsson & Gibson 2004; Gao & Pan 2001; Tatar et al. 2001), genetic background (Chandler et al. 2013; Dworkin & Gibson 2006; Paaby & Rockman 2014) and environment (Ghosh et al. 2013; Shingleton et al. 2009; de Moed et al. 1997). Our results are unique in that they allow us to directly assess the effects of these perturbations on relative growth based on sex for both direction and magnitude. Relative differences between male and female growth patterns due to these mutations are ultimately responsible for the generation of SSD and/or SShD.

### The importance of multiple independent control lineages

As expected, different wild type strains vary in magnitude and direction of effects for SSD and SShD (Figure 1). The Oregon-R wild type background displays greater dimorphism in both size and shape compared to Sam. Implicit in our results is the understanding that genetic background itself has a profound effect on the underlying wild-type growth pathways and all of the downstream consequences this can have.

Somewhat more surprising is that both SSD and SShD appear quite environmentally sensitive (despite the genotypic effects being relatively insensitive based on our previous work). While great care was taken to reduce the effects of microclimactic variation, edge effects, nutritional variation and even genotypic variation, our results demonstrate that size and shape dimorphism remain highly variable (Figure 1).

There always remains the possibility that environmental variation does not entirely account for the wildtype variation observed. For each backcrossed line, a small amount of genetic information surrounding each p-element insertion site is unavoidable, especially during recombination in final cross with mutants and wild-types. This effect is somewhat unlikely, however, due to the fact that these recombination events are rare and affect only single measured individuals. Regardless, such a large amount of variation in trait values within “isogenic” lines is unexpected. Most studies attribute any such variation within genetically (and environmentally) identical lines to stochastic variation in gene expression (Rea et al. 2005; Kirkwood et al. 2005; Raj & van Oudenaarden 2008). Such claims are, however, outside of the scope of our current study.

### Rare sex-limited effects on wing form among mutations in EGFR and TGF-ß signaling

While much is known about the development of wing size and shape (Shingleton et al. 2005; García-Bellido et al. 1994; Weinkove et al. 1999; Day & Lawrence 2000; Prober & Edgar 2000), comparatively little is known about the sex-specific effects of the genes involved (Horabin 2005; Abbott et al. 2010; Gidaszewski et al. 2009). While these mutations represent only a subset of the almost innumerable potential mutations within and among genes, they serve as a lens through which we can view the sex-limited effects of mutations. It is now clear that only a handful of genes associated with growth may be acting in a sex-dependent manner. Indeed, these results call for a further investigation of the formerly understudied sex-effects of growth pathways.

One such study confirms a link between many of the patterning mutations used in the current study and the development of SSD in the wing (Horabin 2005). In her 2005 paper, Horabin demonstrated that components of the sex-determination pathway (specifically, *Sxl)* were responsible for activating size-regulating genes within the Hedgehog signaling pathway. In fact, of the handful of genes to display sex-limited effects on SSD or SShD, a few were associated with this pathway, including: *Omb, dad* and *Dpp* (Horabin 2005; Abu-Shaar & Mann 1998). This does not appear to be coincidence as these are the only mutants in this pathway that we utilized for this study. Since these mutants only represent a subset of those with sex-limiting effects, we cannot assign causality to this pathway. Instead, this demonstrates that sex-limiting effects of genes interact with more complexity than previously understood; no one pathway appears to be acting in a sex-dependent manner to generate shape/size.

Another candidate pathway involved in the generation of SSD is the Insulin and Insulin-like growth factor (IIS)/Target of Rapamycin (TOR) pathway. Evidence suggests that components of this pathway, such as *InR* (Testa et al. 2013; Shingleton et al. 2005) and *foxo*(Carreira et al. 2011) can contribute to SSD and/or SShD.

Further studies, such as Takahashi & Blanckenhorn (2015) have found that most mutations appear to decrease the SSD of wing form. Our data appear to yield an interesting trend for the direction of SSD based on genetic background. Ostensibly, growth-pathway mutants in the Ore wild type background tend to decrease SSD, whereas mutants that affect SSD in Sam tend to increase it. At this point it is impossible to say if this trend is biologically meaningful, but given that Ore has a greater underlying magnitude of SSD (and is already in conflict with Rensch’ s rule), these mutations may be interfering with genetic mechanisms influencing sexual dimorphism in the Ore background.

Our data is somewhat inconsistent with the findings of another previous mutation screen study, namely those of Carreira et al. (2011), wherein the authors found a much greater proportion of random insertion mutations appeared to have sex-specific effects on wing shape. The reasons for this are as of yet unclear, but may reflect methodology, magnitude of mutational effects used or that in the current study all mutations were limited to two signaling pathways. First, our methods allowed us to effectively tease apart the sex-limited interactions of sex for each genotype pair by plotting them in a size-shape space. Second, the authors used a different wild type genetic background than either that were used in this study (Canton-S). It is clear from this study and others (Dworkin & Gibson 2006; Chandler et al. 2013) that genetic background has an appreciable effect on gene function. At least part of the variation in the number of genes affecting wing SSD must necessarily be due to genetic background effects; however, genetic background effects cannot wholly account for the differences observed. Third, we cannot rule out the effects of dominance when discussing the effects of gene function. The genotype of flies in the study by Carreira et al. (2011) was homozygous for all mutants used. Their lines were chosen specifically for their non-lethal homozygous phenotype, whereas mutation used in our study were chosen irrespective of lethality. Because of this, our flies necessarily had to be heterozygous in order to avoid lethality associate with the homozygous phenotype. Perhaps not all loss-of-function mutants within our study were sufficient to alter the phenotype in a sex-limited manner. Finally, because our mutants were deliberately selected based on their association with wing shape morphogenesis, our results are not strictly comparable to those of Carreira et al. (2011).

### Disentangling mutant-phenotype relationships

Our findings suggest that in most cases when a mutational analysis is performed to understand the genetic architecture of SSD or SShD, it is important to assess whether the mutation is only affecting SSD/SShD or whether it is instead demonstrating some degree of sex-biased influence. Many genes may therefore appear to alter SSD/SShD, but are instead only affected by sex as one of several variables of its expression. This may seem like an arbitrary distinction, but it is important if we are to fully understand the genetic underpinnings of complex phenotypes. Many mutants, such as those in the EGFR signaling pathway used here, are either lethal or at least partially ablate development of certain organs as homozygotes, indicating that these genes are necessary for the development of the organ itself. If heterozygotes have sex-specific effects on size or shape, we cannot necessarily conclude that this gene affects SSD or SShD, but rather that the gene is important for formation of an organ and has sex-dependent effects. Only in the case of genes such as *Mafl*, a gene that has been demonstrated to directly effect SSD in *Drosophila* (Rideout et al. 2012), can we conclude that said gene is affecting SSD and not simply acting in a sex-limited manner.

To fully understand the scope of SSD and SShD, one must precisely define what is meant by size and shape. While the definition of size is relatively straightforward to interpret, shape is somewhat more nuanced. For many organs, shape can essentially be broken down into the relative size of component parts of the larger structure (given that all aspects are homologous). For instance, during development in *Drosophila* there are multiple quadrants of the developing wing imaginal disc whose individual sections may grow more or less in relation to the others, thus altering the “shape” of the wing. Mutant phenotypes may manifest as changes to large sections, such as a widening of the entire central portion of the wing *(Ptc)* or they may be subtler in effect, altering the placement of only a single crossvein (cv-2) (Dworkin & Gibson 2006). While these mutants may have local effects on size, such that they alter shape, what is less clear is whether these mutants are affecting size in a localized manner or the actual shape itself.

The effects of each pathway appear relatively consistent despite differences in genetic background. While mutations within the Egfr pathway tended to affect primarily SShD, those in the TGF-β pathway had a more mixed effect (more frequently affecting SSD). This pattern suggests that genetic background may only alter a mutation’s quantitative effect, rather than its qualitative effect.

Ultimately, our results demonstrate the importance of distinguishing between the relative contributions of each mutation to sexual dimorphism for shape, size or both. Of those mutants with sex-limited effects, even fewer exclusively affect either shape or size dimorphism (Fig.2). While some studies have been successful in artificially altering SSD of specific traits through selection (Bird & Schaffer 1972; Emlen et al. 2005; Reeve & Fairbairn 1996), it is unclear whether whole trait size or simply trait shape (e.g. length) has been altered. Our results demonstrate the need to exercise caution when discussing the effect of mutants on size or shape dimorphism.

### Reassessing the assumption of common allometry

One important method for quantifying “shape” changes involves examining allometric relationships, specifically static allometry, which is the relationship among adult individuals between body size and organ size (Huxley 1932; Stern & Emlen 1999). In fact, one of the most obvious way that males and females can differ is through differences in scaling relationships between body parts; these encompass some of the most obvious sources of variation in the natural world (Bonduriansky & Day 2003; Shingleton et al. 2009). By studying the relationship between two traits (e.g. body vs organ size), we can glean important information about the relative growth of traits and, therefore, the underlying mechanisms of differences in the growth of these traits. Consequently, allometry is an important tool for biologists to assess differences in size and shape dimorphism within (and across) a species. Our results support the claim for the importance of studying allometry by demonstrating that, while some mutants may have sex-limited effects on shape and/or size dimorphism (Fig.2), they do not necessarily effect the relationship between trait shape and size (Fig.4). Many mutants cause significant differences in sexual dimorphism of allometry, but do not necessarily alter SSD or SShD. These results may seem counterintuitive, but it is important to remember that, while changes in SSD or SShD may shift the direction of slope of allometry along one or more dimensions (in shape space), this does not necessarily alter the allometric slope itself (Frankino et al. 2005).

Since D’Arcy Thompson (1917) outlined his approach of how relative changes in body and organ size can be mapped out onto Cartesian coordinates to visualize relative growth, the study of allometry and shape have been closely linked. Modern approaches use similar, albeit much more complicated methods to assess changes in relative landmark positions (Sanger et al. 2013; van der Linde & Houle 2009; Abbott et al. 2010). Ostensibly, one of the downfalls of shape analysis is that shape inherently carries information about its underlying relationship to size, despite the fact that geometric morphometric analyses partially separates it from shape (Gidaszewski et al. 2009; Mosimann 1970; Gould 1966; Nevill et al. 1995). More specifically, size itself is a measurement based on some aspect of shape. If size and shape do not scale isometrically (such that unit increase in size is accompanied by an equal increase in shape), then the underlying co-variation will be reflected in estimates of shape that are disproportionately affected by size (Mosimann 1970). This issue is implicit in the geometry of shapes themselves; as absolute size increases, surface area to volume ratios decrease (Gould 1966). This is particularly bad news for studies wishing to analyze induced changes in shape and size, because it means that the degree of independence between these two variables may be difficult to infer. However, by plotting size on shape and using the residuals from this model, Gidaszewski et al. (2009) were able to effectively eliminate the issue of non-independence with size and shape. These residuals represent the total variation in shape that is not due to allometric effects of size.

Allometric patterns of variation across sex and genotype are necessarily more complicated. While it is known that shape (and shape dimorphism) is strongly influenced by its relationship with size, it is not always clear that the assumption of a common allometric relationship across sexes is met. Previous studies examining patterns of SSD and SShD (Gidaszewski et al. 2009) generally made the assumption of a common allometric relationship between males and females within each *Drosophila* species. This was despite their analysis suggesting that this assumption may not hold for all species. For the data we examined here, we could reject this assumption based on inferences based on statistical significance. Yet, it is clear that the magnitude of such differences were small, and allometric relationships were similar in most cases. Indeed, the allometric influence of size on shape appears to be largely consistent with respect to direction of effects, with a few notable exceptions (Fig.4). Regardless, we erred on the side of caution with this matter and decided to eschew analysis of SSD and SShD under assumption of common allometry. It is worth noting that the assumption of common allometry did not substantially alter the observed results (Supplementary Figure 1). As with other studies, we suggest that a rejection of this assumption simply based upon significance may not be optimal, and future work should determine what the consequences of making such assumptions might be for studies of sexual dimorphism and allometry.

Our results clearly demonstrate the effects of growth pathway mutants on SSD and SShD. Most notably, we cannot rule out the sex-specific effects of any genes involved in growth. Our results demonstrate the current lack of understanding of how growth-related genes interact with the sex of the individual. By visualizing the effects of each mutation within the framework of size/shape space we gain a previously unrealized understanding of the role each mutant plays in generating a sex’s phenotype. While this method is especially powerful for studying sexual dimorphism, its applications are not restricted to it. We therefore present this method as a means for dissecting the contributions of mutants to the development of size and shape.

## Acknowledgements

We would like to thank Dr. William Pitchers for providing us with R scripts that facilitated the analysis.

***Funding***: This work was supported by NIH grant 1R01GM094424—01 and an NSERC Discovery award to ID.

***Conflict of Interest***: The authors declare that they have no conflict of interest.

---

**Figure S1.**
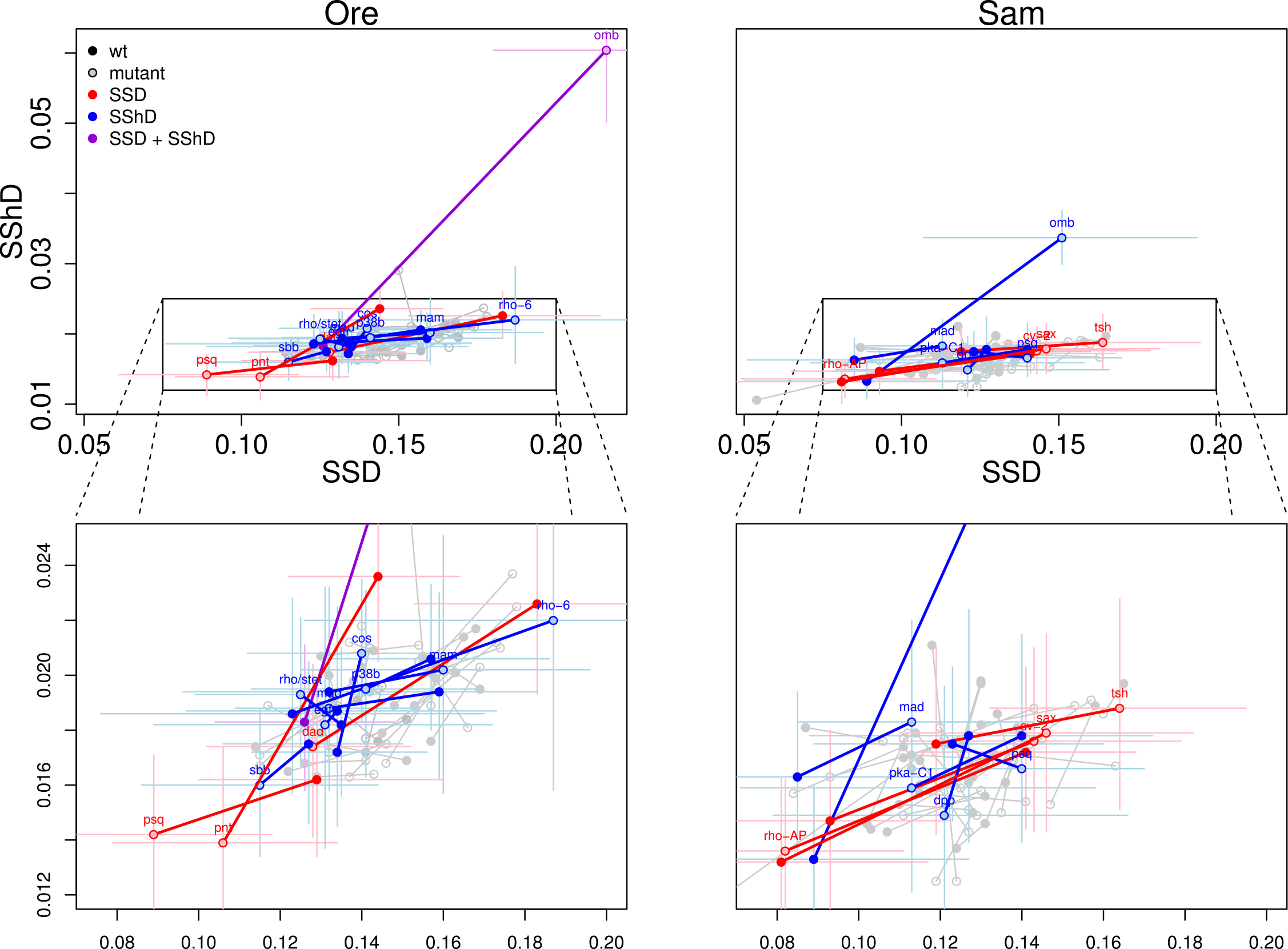
Magnitude of SSD and SShD for 42 mutants in Oregon-R (left) and Samarkand (right) wild type backgrounds, after correcting for the influence of allometry (shape on size). A common allometry relationship was assumed across genotype and sex within each background and line combination. Residuals from the allometric model were then used for the analysis. This figure is otherwise identical to figure 2 (which does not correct for allometry). The effect of each mutant is mapped out in a size-and-shape dimorphism space. Genotypic means for each mutant are indicated by point style and connected by a solid line. SSD is plotted on each x-axis for all plots and SShD is displayed on the y-axis. The plots above display the entire range of variation observed, while those below display only the area with the highest density of points. Lines with significant sex-by-genotype effects are highlighted as follows: effect on both size and shape, shape only and size only. Only significant genes (after sequential Bonferroni correction) from the linear models are colored. Error bars are 95% confidence intervals (unadjusted alpha). All gene names are displayed lower-case, regardless of dominance.

**Supplementary Table 1a.** Significance Table (raw p values) for effects on wing shape after the effects of lometry are removed where G=genotype, S=sex, B=background and Cs=(centroid) size

| Mutant | Line | Allele | Pathway | G | G x S | G x B | Cs x G x S | Cs x G x B | G x S x B | Cs x G x S x B |
| --- | --- | --- | --- | --- | --- | --- | --- | --- | --- | --- |
| *aos* | 2513 | W11 | Egfr | 0.001 | 0.671 | 0.14 | 0.863 | 0.827 | 0.324 | 0.093 |
| *omb* | 3045 | md653 | TGF-β | 0.001 | 0.001 | 0.003 | 0.004 | 0.001 | 0.001 | 0.086 |
| *cv-2* | 6342 | 225-3 | TGF-β | 0.092 | 0.363 | 0.093 | 0.023 | 0.822 | 0.027 | 0.677 |
| *GAP1* | 6372 | mip-w[+] | Egfr | 0.001 | 0.034 | 0.001 | 0.674 | 0.881 | 0.405 | 0.428 |
| *ksr* | 10212 | J5E2 | Egfr | 0.001 | 0.935 | 0.193 | 0.057 | 0.89 | 0.364 | 0.248 |
| *dad* | 10305 | J1E4 | TGF-β | 0.001 | 0.054 | 0.593 | 0.24 | 0.058 | 0.098 | 0.296 |
| *drk* | 10372 | k02401 | Egfr | 0.001 | 0.046 | 0.482 | 0.478 | 0.544 | 0.57 | 0.796 |
| *bs/DSRF* | 10413 | k07909 | Egfr | 0.001 | 0.111 | 0.001 | 0.157 | 0.163 | 0.437 | 0.165 |
| *s* | 10418 | k09530 | Egfr | 0.001 | 0.463 | 0.019 | 0.849 | 0.23 | 0.518 | 0.002 |
| *spi* | 10462 | s3547 | Egfr | 0.001 | 0.518 | 0.003 | 0.495 | 0.52 | 0.438 | 0.573 |
| *mad* | 10474 | k00237 | TGF-β | 0.001 | 0.56 | 0.001 | 0.542 | 0.464 | 0.125 | 0.094 |
| *ed* | 10490 | k01102 | Egfr | 0.001 | 0.88 | 0.012 | 0.07 | 0.051 | 0.542 | 0.046 |
| *tsh* | 10842 | A3-2-66 | TGF-β | 0.095 | 0.458 | 0.397 | 0.947 | 0.775 | 0.165 | 0.262 |
| *cos* | 11156 | k16101 | Hh | 0.011 | 0.228 | 0.001 | 0.267 | 0.096 | 0.288 | 0.36 |
| *tkv* | 11191 | k19713 | TGF-β | 0.001 | 0.121 | 0.001 | 0.358 | 0.813 | 0.26 | 0.161 |
| *babo* | 11207 | k16912 | TGF-β/Hh | 0.054 | 0.428 | 0.047 | 0.465 | 0.51 | 0.429 | 0.1 |
| *trl* | 12088 | S2325 | TGF-β | 0.001 | 0.136 | 0.315 | 0.046 | 0.82 | 0.997 | 0.23 |
| *rho-AP* | 12413 | BG00314 | ? | 0.523 | 0.006 | 0.005 | 0.983 | 0.815 | 0.196 | 0.098 |
| *pka-C1* | 12752 | BG02142 | Hh | 0.225 | 0.946 | 0.356 | 0.032 | 0.89 | 0.825 | 0.601 |
| *sbb* | 12772 | BG01610 | TGF-β | 0.001 | 0.203 | 0.038 | 0.323 | 0.647 | 0.02 | 0.539 |
| *psq* | 12916 | kg00811 | Egfr | 0.011 | 0.38 | 0.223 | 0.979 | 0.732 | 0.03 | 0.01 |
| *osa* | 12945 | kg03117 | Chromatin Remodeling | 0.001 | 0.188 | 0.036 | 0.876 | 0.373 | 0.043 | 0.123 |
| *rasGAP* | 13311 | kg02382 | Egfr | 0.541 | 0.371 | 0.001 | 0.037 | 0.038 | 0.286 | 0.307 |
| *pnt* | 13535 | kg04968 | Egfr | 0.002 | 0.063 | 0.021 | 0.173 | 0.752 | 0.504 | 0.609 |
| *drk* | 13943 | k02401 | Egfr | 0.095 | 0.223 | 0.001 | 0.514 | 0.22 | 0.107 | 0.222 |
| *cbl* | 13944 | kg03080 | Egfr | 0.003 | 0.459 | 0.018 | 0.765 | 0.57 | 0.947 | 0.195 |
| *mam* | 14189 | kg02641 | N/Egfr | 0.001 | 0.908 | 0.577 | 0.298 | 0.035 | 0.245 | 0.066 |
| *rho-6* | 14208 | kg05638 | Egfr | 0.809 | 0.35 | 0.814 | 0.448 | 0.72 | 0.152 | 0.25 |
| *dpp* | 14268 | kg04600 | TGF-β | 0.006 | 0.898 | 0.022 | 0.285 | 0.058 | 0.912 | 0.025 |
| *pka-C3* | 14345 | kg00222 | Hh | 0.177 | 0.045 | 0.035 | 0.587 | 0.276 | 0.1 | 0.018 |
| *p38b* | 14364 | kg01337 | TGF-β/Egfr | 0.247 | 0.59 | 0.031 | 0.566 | 0.906 | 0.771 | 0.902 |
| *tkv* | 14403 | kg01923 | TGF-β | 0.041 | 0.111 | 0.009 | 0.222 | 0.835 | 0.501 | 0.269 |
| *wmd* | 14541 | kg07581 | Unknown | 0.093 | 0.264 | 0.04 | 0.651 | 0.157 | 0.039 | 0.378 |
| *mad* | 14578 | kg00581 | TGF-β | 0.002 | 0.651 | 0.153 | 0.815 | 0.199 | 0.216 | 0.753 |
| *ast* | 14638 | kg07563 | Egfr | 0.014 | 0.972 | 0.074 | 0.211 | 0.934 | 0.666 | 0.781 |
| *dpp* | 14694 | kg08191 | TGF-β | 0.002 | 0.822 | 0.074 | 0.162 | 0.878 | 0.421 | 0.424 |
| *rho1* | 14901 | kg01774 | Egfr? | 0.082 | 0.059 | 0.353 | 0.083 | 0.677 | 0.082 | 0.03 |
| *sax* | 14920 | kg07525 | TGF-β | 0.046 | 0.24 | 0.003 | 0.31 | 0.519 | 0.5 | 0.389 |
| *sax\** | 5404 | sax4 | TGF-β | 0.04 | 0.001 | 0.026 | 0.205 | 0.701 | 0.808 | 0.795 |
| *egfr* | 10385 | k05115 | Egfr | 0.001 | 0.098 | 0.023 | 0.12 | 0.112 | 0.077 | 0.316 |
| *src42A* | 13751 | kg02515 | Egfr | 0.001 | 0.057 | 0.023 | 0.054 | 0.212 | 0.617 | 0.13 |
| *rho/stet* | 14321 | kg07115 | Egfr | 0.001 | 0.653 | 0.305 | 0.064 | 0.558 | 0.102 | 0.327 |

**Supplementary Table 1b.** Sample size of treatments, where M=Male, F=female, W=wild-type m=mutant, O=Oregon-R background, S=Samarkand background

| Mutant | Line | M, w, O | F, w, O | M, w,<br>S | F, w, S | M, m, O | F, m, O | M, m, S | F, m, S |
| --- | --- | --- | --- | --- | --- | --- | --- | --- | --- |
| aos | 2513 | 21 | 20 | 21 | 20 | 20 | 20 | 20 | 22 |
| omb | 3045 | 19 | 18 | 20 | 20 | 12 | 19 | 19 | 20 |
| cv-2 | 6342 | 10 | 11 | 20 | 20 | 10 | 10 | 20 | 20 |
| GAP1 | 6372 | 19 | 18 | 20 | 19 | 18 | 19 | 20 | 20 |
| ksr | 10212 | 17 | 20 | 19 | 20 | 20 | 20 | 22 | 20 |
| dad | 10305 | 20 | 20 | 18 | 20 | 20 | 20 | 20 | 20 |
| drk | 10372 | 20 | 20 | 20 | 20 | 20 | 20 | 20 | 21 |
| bs/DSRF | 10413 | 19 | 19 | 20 | 20 | 20 | 19 | 20 | 20 |
| s | 10418 | 20 | 20 | 18 | 21 | 20 | 20 | 20 | 20 |
| spi | 10462 | 18 | 20 | 20 | 20 | 20 | 19 | 21 | 21 |
| mad | 10474 | 19 | 20 | 20 | 22 | 20 | 20 | 18 | 22 |
| ed | 10490 | 20 | 20 | 19 | 20 | 20 | 20 | 21 | 21 |
| tsh | 10842 | 19 | 21 | 20 | 20 | 20 | 19 | 22 | 19 |
| cos | 11156 | 19 | 20 | 19 | 20 | 22 | 19 | 20 | 18 |
| tkv | 11191 | 20 | 20 | 20 | 21 | 20 | 21 | 18 | 17 |
| babo | 11207 | 19 | 21 | 20 | 20 | 20 | 20 | 20 | 20 |
| trl | 12088 | 20 | 20 | 21 | 20 | 20 | 21 | 20 | 20 |
| rho-AP | 12413 | 20 | 20 | 20 | 20 | 20 | 19 | 20 | 19 |
| pka-C1 | 12752 | 20 | 20 | 18 | 20 | 20 | 20 | 21 | 21 |
| sbb | 12772 | 20 | 20 | 20 | 19 | 20 | 22 | 20 | 21 |
| psq | 12916 | 21 | 20 | 21 | 20 | 20 | 20 | 20 | 21 |
| osa | 12945 | 31 | 30 | 20 | 20 | 30 | 24 | 20 | 20 |
| rasGAP | 13311 | 19 | 19 | 20 | 19 | 21 | 21 | 20 | 20 |
| pnt | 13535 | 20 | 20 | 20 | 19 | 20 | 19 | 20 | 20 |
| drk | 13943 | 20 | 19 | 20 | 21 | 16 | 17 | 21 | 21 |
| cbl | 13944 | 20 | 20 | 21 | 20 | 20 | 20 | 20 | 20 |
| mam | 14189 | 10 | 8 | 21 | 21 | 10 | 10 | 20 | 20 |
| rho-6 | 14208 | 19 | 21 | 20 | 20 | 19 | 20 | 20 | 21 |
| dpp | 14268 | 21 | 20 | 19 | 20 | 20 | 18 | 22 | 18 |
| pka-C3 | 14345 | 21 | 20 | 20 | 20 | 21 | 20 | 20 | 20 |
| p38b | 14364 | 10 | 10 | 20 | 20 | 10 | 6 | 20 | 20 |
| tkv | 14403 | 11 | 10 | 10 | 12 | 10 | 11 | 20 | 20 |
| wmd | 14541 | 20 | 20 | 20 | 21 | 20 | 21 | 21 | 20 |
| mad | 14578 | 20 | 19 | 22 | 21 | 21 | 19 | 20 | 20 |
| ast | 14638 | 20 | 20 | 20 | 20 | 20 | 18 | 20 | 20 |
| dpp | 14694 | 20 | 19 | 19 | 20 | 20 | 20 | 20 | 20 |
| rho1 | 14901 | 20 | 20 | 20 | 21 | 20 | 20 | 20 | 20 |
| sax | 14920 | 19 | 19 | 20 | 20 | 20 | 20 | 21 | 20 |
| sax* | 5404 | 20 | 21 | 18 | 19 | 20 | 20 | 20 | 20 |
| egfr | 10385 | 20 | 19 | 20 | 20 | 20 | 20 | 20 | 20 |
| src42A | 13751 | 19 | 20 | 19 | 20 | 19 | 20 | 19 | 20 |
| rho/stet | 14321 | 19 | 19 | 19 | 20 | 19 | 21 | 19 | 19 |

**Supplementary Table 2.** Significance Table (raw p values) for effects on wing shape after the effects of allometry are removed where G=genotype, S=sex, B=background and Cs=(centroid) size.

| **Mutant** | **Line ID** | **G** | **G x S** | **G x B** | **Cs x G x S** | **Cs x G x B** | **G x S x B** | **Cs x G x S x B** |
| --- | --- | --- | --- | --- | --- | --- | --- | --- |
| *aos* | 2513 | 0.098 | 0.221 | 0.001 | 0.664 | 0.31 | 0.236 | 0.625 |
| *omb* | 3045 | 0.001 | 0.072 | 0.001 | 0.6 | 0.311 | 0.002 | 0.264 |
| *cv-2* | 6342 | 0.066 | 0.215 | 0.171 | 0.494 | 0.412 | 0.619 | 0.208 |
| *GAP1* | 6372 | 0.002 | 0.235 | 0.001 | 0.075 | 0.012 | 0.177 | 0.299 |
| *ksr* | 10212 | 0.008 | 0.005 | 0.002 | 0.165 | 0.24 | 0.165 | 0.526 |
| *dad* | 10305 | 0.16 | 0.009 | 0.797 | 0.158 | 0.003 | 0.962 | 0.205 |
| *drk* | 10372 | 0.005 | 0.174 | 0.031 | 0.115 | 0.021 | 0.021 | 0.607 |
| *bs/DSRF* | 10413 | 0.001 | 0.126 | 0.001 | 0.204 | 0.021 | 0.005 | 0.427 |
| *s* | 10418 | 0.001 | 0.002 | 0.106 | 0.377 | 0.188 | 0.009 | 0.107 |
| *spi* | 10462 | 0.001 | 0.527 | 0.002 | 0.558 | 0.371 | 0.025 | 0.359 |
| *mad* | 10474 | 0.001 | 0.019 | 0.005 | 0.594 | 0.656 | 0.005 | 0.746 |
| *ed* | 10490 | 0.001 | 0.778 | 0.591 | 0.998 | 0.225 | 0.393 | 0.164 |
| *tsh* | 10842 | 0.008 | 0.216 | 0.002 | 0.837 | 0.828 | 0.051 | 0.988 |
| *cos* | 11156 | 0.001 | 0.019 | 0.037 | 0.025 | 0.03 | 0.036 | 0.375 |
| *tkv* | 11191 | 0.001 | 0.004 | 0.006 | 0.404 | 0.947 | 0.301 | 0.135 |
| *babo* | 11207 | 0.048 | 0.427 | 0.02 | 0.208 | 0.517 | 0.721 | 0.092 |
| *trl* | 12088 | 0.001 | 0.001 | 0.04 | 0.776 | 0.891 | 0.005 | 0.38 |
| *rho-AP* | 12413 | 0.075 | 0.444 | 0.017 | 0.805 | 0.69 | 0.194 | 0.009 |
| *pka-C1* | 12752 | 0.001 | 0.227 | 0.107 | 0.081 | 0.513 | 0.002 | 0.204 |
| *sbb* | 12772 | 0.001 | 0.537 | 0.004 | 0.703 | 0.001 | 0.396 | 0.069 |
| *psq* | 12916 | 0.004 | 0.283 | 0.143 | 0.677 | 0.127 | 0.048 | 0.823 |
| *osa* | 12945 | 0.001 | 0.255 | 0.02 | 0.699 | 0.492 | 0.728 | 0.368 |
| *rasGAP* | 13311 | 0.027 | 0.666 | 0.031 | 0.565 | 0.587 | 0.792 | 0.607 |
| *pnt* | 13535 | 0.016 | 0.199 | 0.111 | 0.439 | 0.199 | 0.659 | 0.293 |
| *drk* | 13943 | 0.002 | 0.005 | 0.001 | 0.414 | 0.905 | 0.081 | 0.051 |
| *cbl* | 13944 | 0.138 | 0.07 | 0.329 | 0.148 | 0.002 | 0.754 | 0.518 |
| *mam* | 14189 | 0.001 | 0.15 | 0.043 | 0.001 | 0.109 | 0.043 | 0.386 |
| *rho-6* | 14208 | 0.086 | 0.091 | 0.433 | 0.664 | 0.959 | 0.009 | 0.099 |
| *dpp* | 14268 | 0.017 | 0.322 | 0.08 | 0.798 | 0.109 | 0.871 | 0.822 |
| *pka-C3* | 14345 | 0.019 | 0.024 | 0.567 | 0.151 | 0.237 | 0.345 | 0.23 |
| *p38b* | 14364 | 0.003 | 0.498 | 0.171 | 0.542 | 0.001 | 0.466 | 0.6 |
| *tkv* | 14403 | 0.021 | 0.225 | 0.023 | 0.403 | 0.673 | 0.454 | 0.395 |
| *wmd* | 14541 | 0.028 | 0.118 | 0.435 | 0.248 | 0.599 | 0.001 | 0.172 |
| *mad* | 14578 | 0.001 | 0.038 | 0.026 | 0.639 | 0.456 | 0.357 | 0.11 |
| *ast* | 14638 | 0.078 | 0.091 | 0.002 | 0.995 | 0.317 | 0.007 | 0.132 |
| *dpp* | 14694 | 0.001 | 0.299 | 0.045 | 0.476 | 0.027 | 0.56 | 0.801 |
| *rho1* | 14901 | 0.221 | 0.362 | 0.436 | 0.366 | 0.329 | 0.394 | 0.244 |
| *sax* | 14920 | 0.002 | 0.394 | 0.38 | 0.758 | 0.272 | 0.135 | 0.338 |
| *sax\** | 5404 | 0.007 | 0.007 | 0.047 | 0.442 | 0.929 | 0.299 | 0.845 |
| *egfr* | 10385 | 0.001 | 0.125 | 0.03 | 0.742 | 0.393 | 0.014 | 0.211 |
| *src42A* | 13751 | 0.031 | 0.37 | 0.002 | 0.384 | 0.195 | 0.411 | 0.586 |
| *rho/stet* | 14321 | 0.002 | 0.156 | 0.19 | 0.02 | 0.12 | 0.207 | 0.421 |

